# Post release immune responses of Tasmanian devils vaccinated with an experimental devil facial tumour disease vaccine

**DOI:** 10.1101/2020.12.06.408963

**Authors:** Ruth Pye, Jocelyn Darby, Andrew Flies, Samantha Fox, Scott Carver, Jodie Elmer, Kate Swift, Carolyn Hogg, David Pemberton, Gregory Woods, A. Bruce Lyons

## Abstract

Disease is increasingly becoming a driver of wildlife population declines and extinction risk. Vaccines have been one of the most successful health interventions in human history, but few have been tested for mitigating wildlife disease. The transmissible cancer, devil facial tumour disease (DFTD), triggered the Tasmanian devil’s (*Sarcophilus harrisii*) inclusion on the international endangered species list. Development of a protective DFTD vaccine would provide a valuable management approach for conservation of the species. In 2016, 33 devils from a DFTD-free insurance population were given an experimental DFTD vaccination prior to their release on the north coast of Tasmania. The release site was already home to an incumbent population of devils, including some individuals with DFTD. To determine the efficacy of the vaccination protocol and the longevity of the response it induced, six trapping trips took place at the site over the 2.5 years following release. Eight of the 33 vaccinated devils were re-trapped, and six of those developed DFTD within the monitoring period. Despite the apparent lack of protection provided by the vaccine for the re-trapped devils, we observed several signs of immune activation not usually found in unvaccinated devils. Firstly, sera collected from the eight devils showed that anti-DFTD antibodies persisted for up to two years post vaccination. Secondly, tumour infiltrating lymphocytes were found in three out of four biopsies collected from vaccinated devils which contrasts with the “immune deserts” typical of DFT’s; only one out of twenty incumbent devils with DFTD trapped during the same period had a tumour biopsy exhibiting immune cell infiltrate. Thirdly, immunohistochemical analysis of tumour biopsies from the vaccinated devils identified the functional immune molecules associated with antigen presenting cells (MHC-II) and T cells (CD3), and the immune checkpoint molecule PD-1, all associated with anti-tumour immunity in other species. These results correlate with our previous study on captive devils in which a prophylactic vaccine primed the devil immune system and, following DFTD challenge and tumour growth, immunotherapy induced complete tumour regressions. The field trial results presented here provide further evidence that the devil immune system can be primed to recognise DFTD cells, but additional immune manipulation could be needed for complete protection or induction of tumour regressions.

## INTRODUCTION

Vaccines are increasingly recognised as valuable conservation management tools as emerging diseases threaten wildlife populations, and in some cases entire species (Langwig et al., 2015). Limited attempts have been made to use vaccines for wildlife disease: erysipelas vaccine in kakapo (*Strigops habroptilus*); sylvatic plague vaccine in prairie dogs (*Cynomys spp);* rabies vaccine in Ethiopian wolves (*Canis simensis*) (Gartrell et al., 2005, Rocke et al., 2017, Sillero-Zubiri et al., 2016). Factors that impede the use of safe and protective vaccines for wildlife include the lack of species-specific reagents to test immune responses, the logistics of captive animal and field trials, and the delivery of vaccines to wild populations. The difficulty of addressing wildlife disease with vaccines becomes even more problematic when the vaccine for a novel wildlife disease has to first be developed. Transmissible cancers, such as those seen in the Tasmanian devil (*Sarcophilus harrisii*), are novel pathogens that break many rules of host-pathogen systems, as the infectious agent is a genetically mismatched tumour cell, rather than a more conventional viral or bacterial pathogen.

The Tasmanian devil is the largest extant carnivorous marsupial and unique to Australia’s island state of Tasmania. The species was listed as endangered by the International Union for the Conservation of Nature in 2008 due to devil facial tumour disease (DFTD), a transmissible cancer first observed in 1996 (Hawkins et al., 2006). In 2014 a second independent transmissible cancer was discovered in devils and named DFT2 (Pye et al., 2016b). To date DFT2 has been confirmed only on a peninsula in southeast Tasmania and its future impact on the devil population remains unknown. The first cancer is now commonly referred to as DFT1. DFT1 and DFT2 are two of only three transmissible cancers known to occur in vertebrates worldwide (Murgia et al., 2006, Metzger et al., 2016).

The low immunogenicity of DFT1 is primarily due to downregulation of MHC-I expression on the tumour cell surface (Siddle et al., 2013). MHC-I expression on DFT1 cells is restored in the presence of the inflammatory cytokine interferon gamma (IFN-γ). This mechanism served as the basis of successful captive trials in which healthy devils were immunised with an experimental vaccine prior to being challenged with DFT1 cells. Most of the devils developed DFT1, but subsequent immunotherapy was able to induce DFT1 regressions in half of the devils that had been primed by the vaccine (Tovar et al., 2017). Additionally, a few natural DFT1 regressions have been observed in wild devils, and serological assays have demonstrated that MHC-I is a major target of antibody responses (Pye et al., 2016a, Tovar et al., 2017, Ong et al., 2020). The demonstration of induced DFT1 regressions in captivity, and naturally occurring anti-DFT1 immune responses, suggest that a DFT1 vaccine could be developed for prophylactic use in wild devils.

The initial field immunisation trials took place in 2015 and 2016. These trials were incorporated into the Wild Devil Recovery project, an initiative of the Tasmanian state government’s Save the Tasmanian Devil Program. This ongoing project aims to prevent inbreeding depression and maintain or restore ecosystem function by supplementing devil populations that have been decimated by DFTD (Fox et al., 2019). Fifty-two Tasmanian devils were vaccinated with an experimental DFT1 vaccination protocol and released at two different sites on Tasmania’s north coast: Narawntapu National Park (NNP) and Stony Head (SH). The results of the DFT1 vaccination trials confirmed that nearly all the vaccinated devils produced anti-DFT1 antibodies (Pye et al., 2018). However, evaluation of the efficacy of the vaccinations required long-term monitoring. This was hampered by the modest re-trapping rate of vaccinated devils during the monitoring trips, and the unknown DFT1 exposure the devils received post-release. This was particularly evident at NNP where only two vaccinated devils, both free of DFT1 were re-trapped in the half to two year post-release period. This low re-trapping number could have been due to the higher incidence of vehicle strike of devils released at NNP compared to devils released at SH, and the dispersal of devils beyond the trap lines (Grueber et al., 2017). With the limited long-term data from vaccinated devils at NNP, the focus of this study is the second site, SH, where 33 vaccinated devils were released in August 2016.

The 33 devils were fitted with GPS collars prior to their release at SH, and frequent trapping was carried out at the site over the following four months. The collars were manually removed from all devils within eight weeks of release, and by December 2016, 17 of the devils had received a booster vaccination. In 2017, three of the vaccinated devils were known to have developed DFT1 as a consequence of natural exposure after their release at SH (Pye et al., 2018). While it was clear the anti-DFT1 response induced by the vaccinations was not completely protective, it did result in a shift from the “immune deserts” of typical DFT1 biopsies which have scant, if any, immune cell infiltrate (Loh et al., 2006) to a tumour microenvironment containing MHC-II+ antigen presenting cells, CD3+ T cells, and cells with the immune activation marker PD-1.

## MATERIALS AND METHODS

### Field trips

Thirty-three vaccinated devils, all permanently marked with a microchip transponder (Allflex®, Palmerston North, New Zealand) were released at Stony Head in August 2016 (Pye et al., 2018). Samples were collected from six monitoring trips which took place between June 2017 and March 2019. Each trip consisted of seven consecutive trapping days, and 40 traps were set in the same locations on each trip, covering a 50 km^2^ area. Traps were checked daily. The field trip methods were as described in (Lazenby et al., 2018).

On each trip, samples taken from the vaccinated and incumbent devils were: up to 5 ml of blood collected from the jugular vein with a 21G hypodermic needle and 5 ml syringe, and placed in clot activating tubes (MTS, Sarstedt) for subsequent serum separation; fine needle aspirates (FNA) of tumours collected with a 21G hypodermic needle and 5 ml syringe and suspended in cell culture medium; tumour biopsies collected with a 3 mm or 4 mm biopsy punch (Kai Medical) and placed in 10% neutral buffered formalin.

### Karyotype of DFT1 cells

Tumour cells collected from DFT’s by fine needle aspirate were karyotyped according to Pye et al., 2016a and briefly described here. Within 48 hours of collection, the tumour cells were transferred to 25 cm^2^ culture flasks with RPMI 1640 (Sigma-Aldrich) and 10% foetal calf serum (Sigma Aldrich). Mitosis inhibition and fixation steps were carried out over the following days. G banding was conducted by treating slides with trypsin, staining with Leishmann’s stain and mounting with Leica mounting medium (Leica Microsystems) for analysis. Karyotypes were made using VideoTest Karyo 3.1 (VideoTest). Fifty metaphases were analysed for each sample.

### DFT1 biopsies, immunohistochemistry and analysis

Tumour biopsies were collected from all devils with ulcerated DFT1 trapped during the monitoring trips, unless tumours were located on the hard palate of the mouth precluding biopsy. Biopsies were fixed in 10% neutral buffered formalin. 3 μm paraffin sections were cut, and standard haematoxylin and eosin (H&E) and immunohistochemical staining carried out as described in detail in (Tovar et al., 2011). The primary antibodies used for immunohistochemistry were: antihuman HLA-DR (referred to from here as MHC-II) (Dako M0746; 1:40); anti-human CD3 (Dako A0452; 1:300); anti-human periaxin (PXN) (Sigma Aldrich HPA001868; 1:300); anti-devil PD-1 clone 3G8 (neat supernatant); and anti-devil PD-L1 clone 1F8 (neat supernatant) (Flies et al., 2016).

Sections of all the biopsies from vaccinated devils and a representative selection of biopsies from eight incumbent devils were then double stained using PXN and MHC-II; PXN and CD3; and PXN and anti-devil PD-1. In brief: Sections were stained for MHC-II, CD3 or PD-1. After the signal detection DAB staining step, slides were washed with water then PBS. If the primary antibody was rabbit then antigen retrieval was repeated to remove it. Slides were incubated with serum free blocking solution (Dako X0909) then incubated with PXN primary antibody. Signal detection of PXN was performed using polyclonal goat anti-rabbit AP (Dako D0487) and liquid permanent red (Dako K0640). After washing in water, slides were counterstained with haematoxylin. Sections from the vaccinated devils’ and the eight incumbent devils’ biopsies were also (single) stained with PD-L1 according to methods described in (Flies et al., 2016). Tumour infiltrating cells staining positive for MHC-II, CD3, or PD-1 on the double stained sections were counted according to the method described in (Zhang et al., 2003) i.e. the number of infiltrating cells in up to 10 high power fields was counted and the average taken. Prior to statistical analysis, data was examined for normality, and log10 transformed to normalise. Unpaired t-tests were used to compare MHC-II, CD3, and PD-1 expression between vaccinated and incumbent cohorts. If the devils had more than one tumour, or were trapped with DFT1 on more than one occasion, only one tumour on the final trapping date was included in the analysis.

The H&E stained sections were examined by veterinary pathologists. Since biopsies were only 3-4 mm in size, only malignant characteristics relating to cytoplasmic or nuclear atypia were assessed. Other histological characteristics of malignancy such as patterns of growth, and presence or absence of necrosis could not be assessed.

### Serum anti-DFT1 antibody analysis

Serum samples from all vaccinated devils and 76 incumbent devils (**Table S7**) were analysed for serum antibodies against whole DFT1 cells not expressing surface MHC-I, and whole DFT cells treated with IFN-g to generate surface expression of MHC-I. The method incorporated indirect immunofluorescence and flow cytometry and is described in detail in (Pye et al., 2016a). In brief, cultured DFT1 cells (cell line C5065) were treated with recombinant devil IFN-g (5 ng/mL) to induce cell surface expression of MHC-I, confirmed by flow cytometry with positive staining for Beta 2 microglobulin (B2M). These cells are referred to as MHC-I^+ve^ DFT1 cells. Untreated cells, negative for B2M are referred to as MHC-I^-ve^ DFT1 cells. Serum samples diluted 1:50 with flow cytometry buffer were incubated with MHC-I^+ve^ DFT1 cells, and separately with MHC-I^-ve^ DFT1 cells. After washing, cells were incubated with a monoclonal mouse anti-devil IgG antibody. Cells were washed then incubated with a fluorochrome labelled goat anti-mouse IgG antibody (Alexa Fluor®647). After a final wash, cells were resuspended in flow cytometry buffer containing cell viability dye to allow gating of dead cells, and analysed by flow cytometry (BD Canto II) and FlowJO (DeNovo software).

The median fluorescence intensity ratio (MFIR) was used to classify the antibody responses. The MFIR is defined as the median fluorescence intensity (MFI) of DFT1 cells labelled with immune serum divided by the MFI of DFT1 cells labelled with pre-immune serum. This ratio accounts for any background serum IgG present prior to the immunizations and standardizes the responses between individual devils. Antibody responses were classified as negative (MFIR < 1.5), or positive (MFIR ≥ 1.5) (Pye et al., 2018).

## RESULTS

### Field trips

In the 0.5 to 2.5 year post-release period, eight of the 33 vaccinated devils were trapped on one or multiple trips **(Tables 1, 2).** Of the eight devils, two were healthy and six had DFT1 by the end of the 2.5 year period. Four of the six devils with DFT1 had their tumour biopsies collected into formalin. Of the other two devils, one had a tumour on the hard palate and a non-ulcerated tumour precluding biopsy, and the other devil was found dead with autolysis precluding histology.

**Table 1.** The number, sex and DFT1 status of vaccinated devils and incumbent devils trapped during each post release monitoring trip. The number of subadult incumbent devils trapped is also shown.

| Trip date | Vaccinated devils |  |  |  | Incumbent devils |  |  |  |  |
| --- | --- | --- | --- | --- | --- | --- | --- | --- | --- |
|  | Number trapped | Male | Female | DFT1 | Number trapped | Male | Female | Subadults < 2 years old | DFT1 |
| **June 2017** | 5 | 3 | 2 | 3 | 25 | 11 | 14 | 21 | 2 |
| **Sep 2017** | 5 | 4 | 1 | 3 | 22 | 7 | 15 | 19 | 2 |
| **Feb 2018** | 1 | 0 | 1 | 1 | 45 | 21 | 24 | 42 | 5 |
| **May 2018** | 0 | 0 | 0 | 0 | 46 | 18 | 28 | 32 | 8 |
| **Aug 2018** | 1<br>(+1 found dead) | 0 | 2 | 2 | 43 | 19 | 24 | 36 | 5 |
| **March 2019** | 0 | 0 | 0 | 0 | 38 | 20 | 18 | 33 | 2 |
| **Total, without re-traps** | 8 | 5 | 3 | 6 | 124 | 60 | 64 | 103 | 20 |

**Table 2.** The devil identification, sex, year of birth; and vaccination booster and DFT1 status of the individual vaccinated devils (Va1 to Va8) trapped during each post release monitoring trip.

| Devil ID,<br>Sex,<br>YOB | Booster<br>Dec<br>2016 | June 2017 | Sep 2017 | Feb 2018 | May<br>2018 | Aug 2018 | March<br>2019 |
| --- | --- | --- | --- | --- | --- | --- | --- |
| Va1<br>Male,<br>2013 | yes | healthy | healthy | / | / | / | / |
| Va2<br>Male,<br>2014 | no | DFT1<br>T1*: S, U<br>T2: S, NU<br>TTV:NA | / | / | / | / | / |
| Va3<br>Male,<br>2014 | yes | / | healthy | / | / | / | / |
| Va4<br>Female,<br>2013 | no | / | / | / | / | DFT1<br>T1: S, U<br>TTV:2,314 | / |
| Va5<br>Male,<br>2013 | yes | / | DFT1<br>T1: L, U<br>TTV:<br>97,619<br>T2: S, U<br>TTV:<br>4,121 | / | / | / | / |
| Va6<br>Male,<br>2014 | yes | DFT1<br>T1: S, U<br>TTV:283 | DFT1<br>T1: S, U<br>TTV:963<br>T2: S, NU<br>TTV 359<br>T3:S, NU<br>TTV:1,964 | / | / | / | / |
| Va7<br>Female,<br>2012 | yes | DFT1<br>T1: S, U<br>TTV:749 | DFT1<br>T1: S, U<br>TTV:<br>1,152<br>T2*:S, NU<br>TTV:48 | DFT1<br>T1:S, U<br>TTV: 933<br>T2*:S, U<br>TTV: 377<br>T3: S, U<br>TTV: 184<br>T4: S, NU<br>TTV: 762 | / | / | / |
| Va8<br>Female,<br>2014 | yes | healthy | / | / | / | Found<br>dead with<br>DFT1<br>** | / |

YOB: year of birth; / : not trapped; Tumour size: S = small (TTV <5,000mm<sup>3</sup>), M = medium (TTV 5,000 – 35,000mm<sup>3</sup>), L = large (>35,000mm<sup>3</sup>), TTV: total tumour volume (Peck et al., 2016); Tumour appearance: U = ulcerated; NU = non-ulcerated; NA: not available
\* location of tumour on hard palate precluding biopsy ; \*\* autolysis precluding histology

A total of 124 incumbent devils were trapped over the same period **(Table 1)** with 60 of these trapped on multiple trips. Most of the incumbents were subadults (less than two years old) which is a typical age structure for devils in long-diseased areas, but ages ranged from less than one year to five years old. There were 20 individuals that acquired DFT1 at some point over the 2.5 year period **(Table 1)**, and 13 of these devils were adults (two years or older). Of the nine adults that were healthy on the last trip they were trapped and examined, none were more than two years old except for one female who was five years old.

### Karyotype of DFT1 cells

Fine needle aspirates of tumours from 10 different devils (three vaccinated and seven incumbents) grew in cell culture which allowed for karyotyping. All of these were tetraploid strain 1. This is the same strain that was found at Stony Head in 2015. These cells did not proliferate in culture and cell lines could not be established.

### DFT1 biopsies, immunohistochemistry and analysis

Tumour biopsies from four vaccinated and the 20 incumbent devils with DFT1 were assessed. H&E stained sections were examined by veterinary pathologists to confirm the DFT1 diagnoses. Tumours were noted to have mild to moderate anisokaryosis and karyomegaly, and most tumours had 1-2 mitotic figures per high power field. Histopathology findings of the tumour biopsies from the individual devils referred to in the manuscript are summarised in **Table S1**.

All of the formalin fixed tumour biopsies were stained with periaxin (PXN), MHC-II and CD3 antibodies to identify DFT1 cells, antigen presenting cells and T lymphocytes respectively. DFT1 biopsies collected from the four vaccinated devils, and eight of the 20 incumbent devils were then double stained with PXN and CD3; PXN and MHC-II; and PXN and PD-1 antibodies to more readily identify the tumour infiltrating cells (**Figure 1)**. The eight incumbents were selected because the tumour biopsies were representative of most incumbent biopsies (n=7), or because the tumour biopsy was noted to have more than the typical number of MHC-II and CD3 positive cells on the single staining sections (n=1). The eight vaccinated devils are referred to as Va1 to Va8, and the eight incumbent devils as In1 to In8 (**Table S2**).

**Figure 1.**
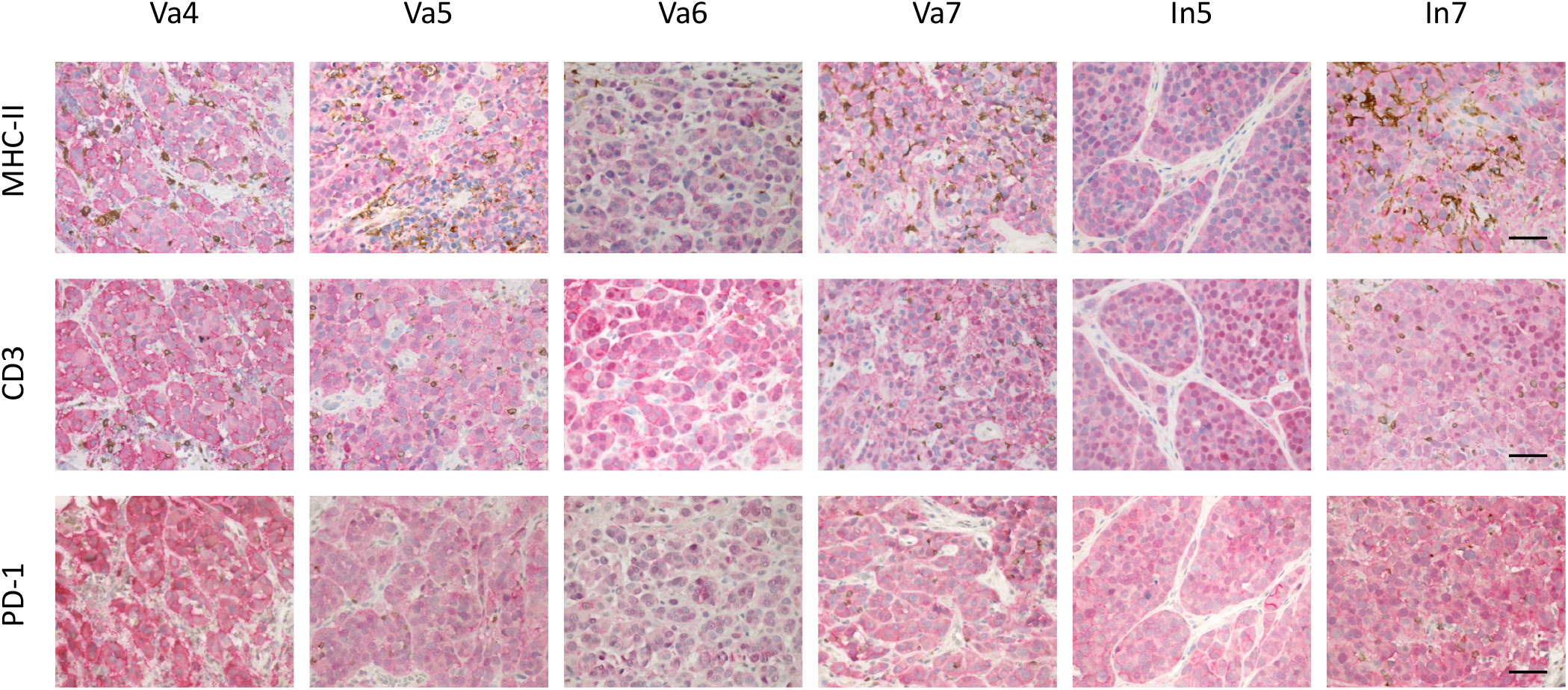
Immunohistochemistry showing immune cell infiltration in DFT1 biopsies from vaccinated and incumbent devils. The individual vaccinated devils are identified as Va4, Va5, Va6 & Va7; and the incumbent devils are In5 & In7. The biopsy from In5 was representative of all the incumbent devils’ tumour biopsies except for In7, which had more tumour infiltrating cells. All images are of immunohistochemical double staining using anti-periaxin antibody to detect DFT1 cells (pink) with either anti-MHC-II (brown), anti-CD3 (brown) or anti-PD-1 (brown) to identify tumour infiltrating cells. Haematoxylin is the counterstain. Scale bar = 50 μm

Biopsies from three of the four vaccinated devils (Va 4, Va5, Va7) had noticeably more tumour infiltrating immune cells staining positive for CD3, MHC-II and PD-1 than biopsies from the incumbent devils **(Figures 1,2, Tables 3, 4, S3, S4, S5)**. One of the incumbent devils (In7) had markedly more tumour infiltrating cells than the other incumbents. This was the only incumbent devil with DFT1 to be trapped and biopsied on consecutive trips. In7 was trapped a third time and although the tumour had increased in size, it was not ulcerated so a biopsy was not collected. The immunohistochemistry results of the three devils (two vaccinated and one incumbent) trapped on consecutive trips are summarised in **Tables S4, S5.** The number of tumour infiltrating cells increased over time, although only slightly for Va6. PD-L1 was rarely expressed, if at all, in the tumour biopsies regardless of immunisation status. Three devils (Va5, In3 and In8) had up to three PD-L1 positive cells in the whole biopsy, including the interstitium (results not shown).

**Figure 2.**
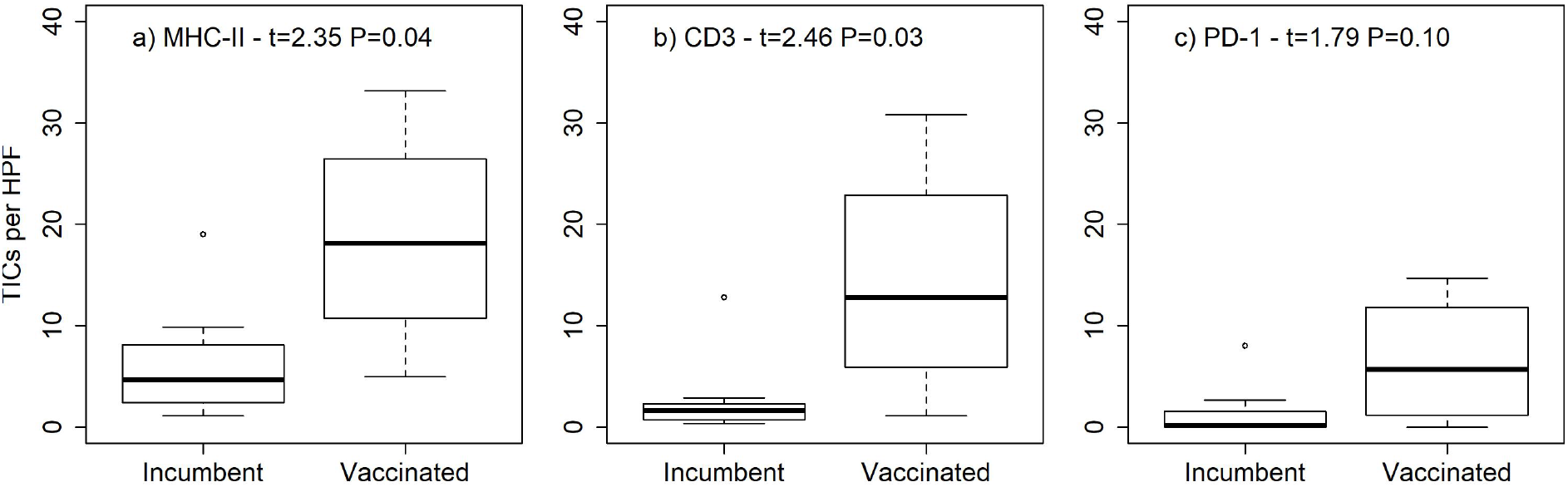
Box and whisker plots comparing the number of tumor infiltrating cells staining positive for a) MHC-II, b) CD3, and c) PD-1 in DFT1 biopsies from vaccinated and incumbent devils. The y axis for each plot is the number of tumor infiltrating cells (TICs) staining positive for each marker per high power field (HPF).

**Table 3.** Summary of the immunohistochemistry analysis of the vaccinated devils’ DFT1 biopsies, showing the number of infiltrating immune cells staining positive for MHC-II, CD3 and PD-1. Vaccinated devils, Va1 to Va8.

| Devil and tumour details |  |  |  | Immunohistochemistry:<br>Number of tumour<br>infiltrating cells staining<br>positive for MHC-II, CD3 or<br>PD-1 |  |  |
| --- | --- | --- | --- | --- | --- | --- |
| Devil ID | Sex, age (in years) at time of sample collection | DFT1 status, date of DFT1 diagnosis (or, if healthy, date last trapped) | Tumour, and date of biopsy if different from date of diagnosis | MHC-II | CD3 | PD-1 |
| Va1 | Male, 4y | Healthy 09/17 | NA | NA | NA | NA |
| Va2 | Male, 3y | DFT1 07/17 | T1, T2 | / | / | / |
| Va3 | Male, 3y | Healthy 09/17 | NA | NA | NA | NA |
| Va4 | Female, 5y | DFT1 08/18 | T1 | ++ | ++ | ++ |
| Va5 | Male, 4y | DFT1 09/17 | T1 | +++ | ++ | + |
|  |  |  | T2 | ++ | ++ | + |
| Va6 | Male, 3y | DFT1 07/17 | T1 09/17 | + | + | 0 |
| Va7 | Female, 5y | DFT1 07/17 | T1 02/18 | +++ | +++ | ++ |
|  |  |  | T3 02/18 | ++ | ++ | + |
| Va8 | Female, 3y | DFT1 Trapped healthy 07/17. Found dead with DFT1 08/18 | T1 | / | / | / |

NA: not applicable
/ : indicates tumour not biopsied either because tumour located on hard palate; not ulcerated; or autolysis
Intratumoural immune cells graded as 0, +, ++, or +++ (0, 1-5, 6 to 19, or $\geq 20$ T cells per high-power field, respectively) (Zhang et al., 2003)
Devils trapped on multiple occasions (i.e. Va6 and Va7) have results shown for the final sampling date

**Table 4.** Summary of the immunohistochemistry analysis of the incumbent devils’ DFT1 biopsies, showing the number of infiltrating immune cells staining positive for MHC-II, CD3 and PD-1. Incumbent devils, In1 to In8.

| Devil and tumour details |  |  |  | Immunohistochemistry:<br>Number of tumour<br>infiltrating cells staining<br>positive for MHC-II, CD3 or<br>PD-1 |  |  |
| --- | --- | --- | --- | --- | --- | --- |
| Devil<br>ID | Sex,<br>Age (in<br>years) at<br>time of<br>sample<br>collection | DFT1<br>status, date<br>of DFT1<br>diagnosis | Tumour<br>biopsied, and<br>biopsy date if<br>different from<br>date of<br>diagnosis | MHC-II | CD3 | PD-1 |
| In1 | Male, 3y | DFT1<br>02/18 | T2 (of 7) | + | + | 0 |
| In2 | Male, 2y | DFT1<br>05/18 | T1 (of 4) | + | + | 0 |
| In3 | Female, 2y | DFT1<br>05/18 | T1 | + | 0 | 0 |
| In4 | Male, 2y | DFT1<br>05/18 | T1 | + | + | 0 |
| In5 | Male, 2y | DFT1<br>05/18 | T1 (of 2) | + | 0 | 0 |
| In6 | Male, 3y | DFT1<br>02/18 | T2 (of 3) | ++ | + | + |
| In7 | Female, 2y | DFT1<br>02/18 | T1<br>05/18 | ++ | ++ | ++ |
| In8 | Female, 3y | DFT1<br>02/18 | T1 | ++ | + | 0 |

Intratumoural immune cells graded as 0, +, ++, or +++ (0, 1-5, 6 to 19, or $\geq 20$ T cells per high-power field, respectively) (Zhang et al., 2003)
In7 was trapped on multiple occasions, the result is shown for the final sampling date

### Serum anti-DFT1 antibody analysis

All the serum samples collected from the eight vaccinated devils on the last date they were re-trapped were positive for IgG antibodies against DFT1 cells **(Figure 3, Table S6)**. Similarly, most of the devils that had a booster vaccination in December 2016 responded with increased serum antibody. Va4 only had a positive antibody response against MHC-I^+ve^ cells **(Figure 3a, Table S6)**, but this devil was the last to be trapped, two years after the vaccination course was completed. Va4 had a minimal antibody response in August 2016 compared to the pre-immune sample **(Figure 3)**. This is explained by the different initial vaccination protocol Va4 received due to not being trapped on the day the first vaccinations were given. The other devils completed their protocol one month prior to release, and serum samples were collected for antibody analysis at the time of release in August. Va4, however, completed the protocol in August, and therefore the peak antibody response, occurring within four weeks of completion, could not be measured for Va4 before release as it was for the other devils (Pye et al., 2018). Va4 and one other (Va2) did not have a booster vaccination in December 2016 which might also account for the lower final antibody response **(Figure 3a, Table 2).** Of the serum samples collected and analysed from 76 incumbent devils (14 with DFT1), three were positive for serum antibodies against DFT1 cells (**Table S6**). One of these devils was a three year old male (In1) with DFT1 and antibodies against both MHC-I^+ve^ and MHC-I^-ve^ DFT1 cells. The other two devils, a three year old female (In8) with DFT1, and a five year old healthy female (In9), had antibodies against MHC-I^+ve^ DFT1 cells only.

**Figure 3.**
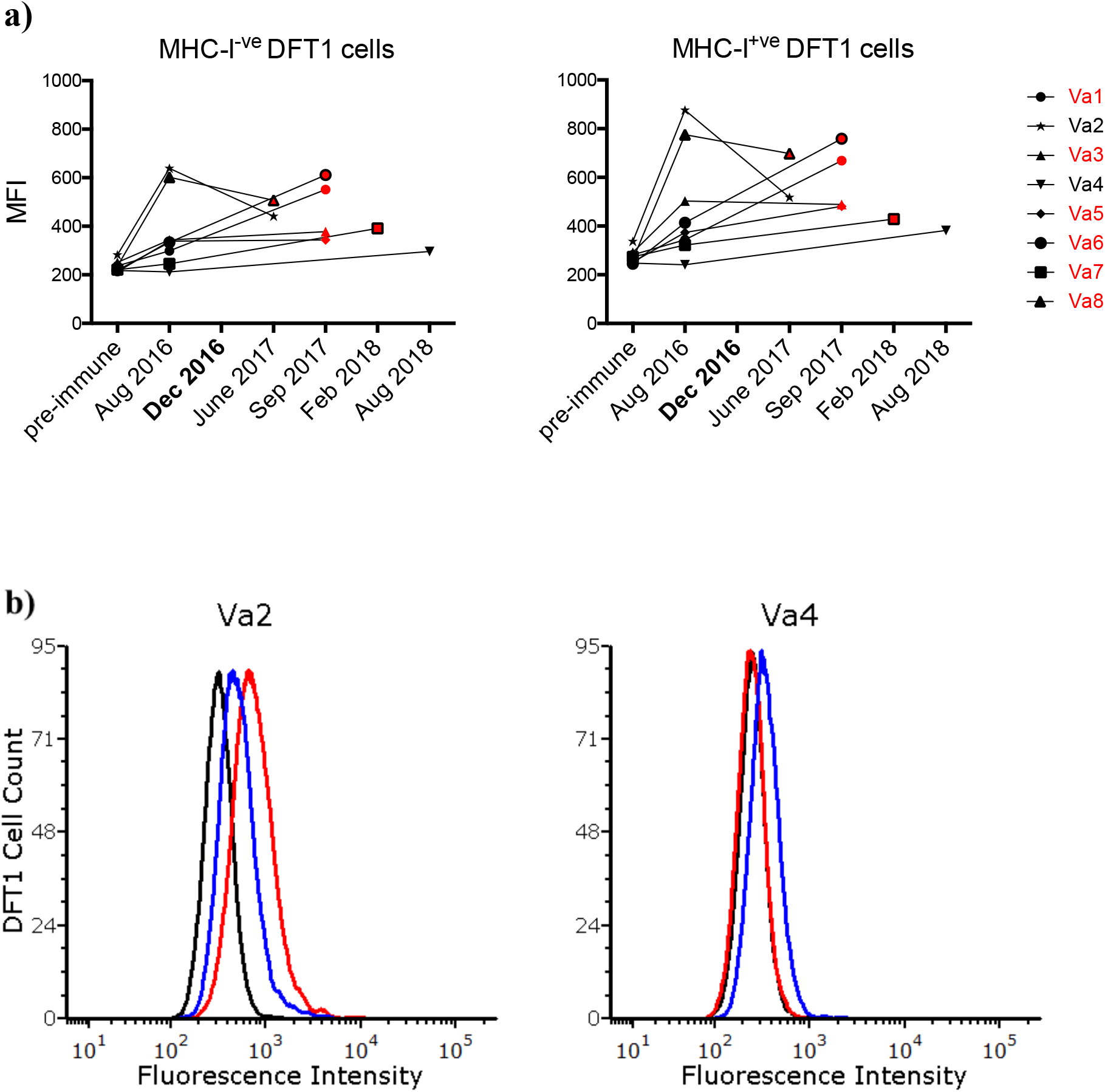
Serum anti-DFT1 IgG antibody responses in vaccinated devils against DFT1 cells. **a)** The graphs show the antibody responses against MHC-I^-ve^ and MHC-I^+ve^ DFT1 cells of the eight vaccinated devils trapped on the monitoring trips. The devils listed in red in the legend had a booster vaccination in December 2016, and the red symbols on the graph indicate their post-booster responses. The y axes show the median fluorescence intensity (MFI) of the serum samples, and the x axes show the dates that serum samples were collected. **b)** The flow cytometry plots are of the serum samples from the devils with the highest (Va2) and the lowest (Va4) antibody responses against DFT1 MHC-I^+ve^ cells. The pre-immune, release and postrelease serum samples are represented by the black, red and blue lines respectively. The y axes show the DFT1 cell count, the x axes show the fluorescence intensity.

## DISCUSSION

This study was a follow-up to the 2015-2016 vaccination field trials that demonstrated a majority of Tasmanian devils could mount an anti-DFT1 immune response (Pye et al., 2018). Modest re-trapping success and the unknown DFT1 exposure of the vaccinated, released devils limited our capacity to determine the level of protection provided by the vaccinations. Despite this, serum and tumour samples collected during the post-release monitoring trips yielded useful insights to the effects of vaccinations on a natural DFT1 challenge.

Eight of the 33 devils released at Stony Head (SH) were trapped during the 0.5 to 2.5 year post release monitoring period. Immediately following their release at SH, the devils’ movements were tracked via their GPS collars until the collars were removed between two and eight weeks later. The devils were found to travel up to 26 km in a single day indicating that many of the devils may not have remained in the immediate release, and post-release trapping area (Samantha Fox, personal communication). The modest re-trapping success of this study could at least be partially explained by these large travel distances.

Six of the eight devils developed DFT1 following natural exposure to the disease, demonstration the vaccine was not protective. The tumour biopsies collected from these devils however, provided evidence of an immune response against the tumour and this is in stark contrast to the immune deserts of typical DFT1 biopsies. Three of the four tumour biopsies (75%) collected from vaccinated devils with DFT1 revealed more tumour infiltrating cells staining positive for MHC-II, CD3 and PD-1 than tumour biopsies from incumbent devils, of which only one out of 20 (5%) had immune cell infiltrate. The low number of tumours from vaccinated devils that could be analysed is an obvious limitation of this study but even so, the results indicate the vaccination was associated with turning what is typically a “cold”, immune-unreactive tumour toward a “hot” tumour. The T cell infiltrate associated with hot tumours results in increased tumour cell mutational load, increased neoantigen presentation and increased immune recognition (Duan et al., 2020). The study on captive devils that underwent immune-mediated regression of experimentally induced DFTs demonstrated that remission correlated with MHC-II and CD3 positive cell infiltration in the tumour biopsies (Tovar et al., 2017). Infiltration of MHC-II positive cells suggests immune recognition, but with the limited panel of antibodies for immunohistochemistry, we cannot infer if these antigen presenting cells are supporting an ongoing anti-tumour response or contributing to immunosuppression. For example, MHC-II positive tumour-associated macrophages are amongst the major tumour-promoting cells found in the tumour microenvironment (Liu et al., 2015).

All eight of the vaccinated devils remained positive for serum anti-DFT1 antibody for up to two years but it is clear that serum antibodies in these devils did not prevent disease. The presence of anti-DFT1 antibodies signifies a response to the vaccination, and the infiltrating T cells found in the biopsies of the vaccinated devils could be a consequence of these antibodies. Antibody dependent cell mediated cytotoxicity (ADCC) might have been triggered in vaccinated devils when challenged with DFT1. In ADCC, antibodies bind to tumour surface antigens and tumour cells can be killed following cross-linking of Fc receptors on effector cells, such as natural killer cells. Natural killer cell activation should also result in secretion of IFN-g to upregulate MHC-I on the DFT cell surface and draw T lymphocytes (CD3 positive) into the tumour. Many other regulatory pathways are involved in the response that can steer the immune system toward further activation or immunosuppression. Since immune deserts are the least immunogenic type of tumours, stimulating immune recognition is an important first step in developing a protective antitumour memory response.

The presence of antibodies in vaccinated devils was likely due to the vaccination, but a contributing effect of subsequent DFT1 exposure cannot be ruled out. There is currently no preclinical diagnostic test for DFTD. Also, while it is possible to distinguish between serum antibodies induced by vaccination against a disease, and antibodies that result from natural exposure to the disease (Westman et al., 2015, Khan et al., 2016), a reliable method to determine this for DFTD is not yet available. Regardless, the longevity of the immune response is encouraging for future DFT1 vaccine development.

Since a high proportion of the re-trapped vaccinated devils developed DFT1 over the two years post-release, it is important to demonstrate that immunisation did not cause, or increase susceptibility to DFT1. The karyotype of the tumours from the seven incumbent and three vaccinated devils was the same tetraploid strain 1. This differs to the diploid strain 3 karyotype of the cell line used in the vaccinations (Tovar et al., 2017) showing the tumours were not derived from the vaccinations. The age of the vaccinated devils may have increased their susceptibility to natural DFT1 challenges. Thirty-two of the 33 devils would have been three years or older in 2017, and older individuals are known to be the first in a population to succumb to DFTD (Lachish et al., 2007).

As discussed above, most of the vaccinated devils that acquired DFT1 and had tumour biopsies collected demonstrated immune recognition of the tumours in the form of increased numbers of MHC-II and CD3 positive infiltrating cells. This study also included PD-1 and PD-L1 in the immunohistochemistry analysis. PD-1 is an immune checkpoint molecule, present on activated T cells (Sharpe et al., 2018). Along with being a marker of effector T cells, PD-1 also plays an essential role in down-regulating the immune response to protect against autoimmunity. PD-1 can also limit protective immunity in the face of chronic disease. The presence of PD-1 positive infiltrating cells in the DFT biopsies correlated with the presence of CD3 and MHC-II positive infiltrating cells. While it is encouraging to think the infiltrating T cells (CD3) were initially activated, the PD-1 expression identified in this study could signify the T cell tolerance which occurs with chronic stimulation. PD-L1 is a transmembrane protein and the ligand for PD-1. Expression of PD-L1 on tumour cells is a powerful immune escape mechanism employed by a variety of different cancers (Azuma et al., 2008). The lack of PD-L1 positive cells in the SH tumour biopsies was in keeping with the 2016 study which found that PD-L1 was generally not expressed in DFT1 aside from the occasional cell (Flies et al., 2016).

Immune responses to DFT1 have previously been reported in a small number of wild devils (Pye et al., 2016a). It was therefore not surprising to find a few incumbent devils with immune responses. There was no obvious pattern amongst these four devils: the two devils with DFT1 and serum antibodies had no infiltrating lymphocytes in their biopsies, and the devil with immune cell infiltrate in the biopsy was negative for serum antibody. The five year old healthy devil with antibody against MHC-I^+ve^ cells is reminiscent of the devils referred to in (Pye et al., 2016a) that had undergone tumour regression. Most of these examples do not fit with the ADCC explanation proposed above for the vaccinated devils in which both serum antibodies and tumour infiltrating cells were present. It is plausible that a number of mechanisms can result in immune responses against DFT1, with differing outcomes. The presence of antibodies only against MHC-I^+ve^ DFT cells in some devils raises the likelihood that MHC-I type of individual devils could play a role in immune recognition. There is already evidence that MHC-I on DFT cells is a target for anti-tumour immune responses in some devils with naturally occurring DFT1 regression (Ong et al., 2020). Deep sequencing of the MHC-I of devils with these responses might reveal the significance of MHC-I type as it relates to DFT1 infection.

The incumbent devil (In7) with immune cell infiltration in the tumour biopsy was trapped on three separate occasions with DFT1, as was only one other devil in the study, Va7, one of the vaccinated cohort. The tumours from these two devils had the highest immune cell infiltrate of all the tumours analysed, and given they were the only devils with DFT1 trapped on more than two separate trips, it is likely their survival time was longer than the other devils with DFT1. Notably, the size of the tumours in both these devils remained “small” over the time they were trapped. It is possible that the immune cell infiltration, induced in Va7 and naturally occurring in In7, played a role in the slow tumour growth rate and increased survival time. Immune cell infiltration slowing tumour growth and/ or increasing survival time has been documented in humans (Berghoff et al., 2016, Zhang et al., 2003). For DFT1, this outcome might be a two-edged sword: improved survival time of a devil with DFT1 would likely improve breeding outcomes for the individual but simultaneously increase the probability of tumour transmission due to a prolonged infectious period.

To date, DFT1 vaccine candidates have been based on whole tumour cells, either irradiated cells or cell lysates from the cultured cell line. This cell line has some karyotypic differences to the wild type DFT1 tumours, raising questions of its appropriateness as a vaccine base. However, recent research on DFT1 evolution shows the tumour to be remarkably stable and, although some DNA copy number differences are found between DFT1 biopsies and cell lines, these invariably involve just one of the two chromosomal copies (Kwon et al., 2020). Cell lines should therefore express similar tumour antigens as wild type tumours. As there were limitations with the whole DFT1 cell vaccine, alternative vaccine approaches such as the use of oncolytic viral vectors that express tumour-associated antigens are worth pursuing (Flies et al., 2020b).

Vaccination for controlling wildlife disease can be a useful conservation tool. Where existing vaccines are available, difficulties lie in their delivery to wildlife, and the assessment of immune responses. Oral bait vaccine platforms go a long way toward addressing vaccine distribution and delivery (Flies et al., 2020b). Similarly, a recently published method to produce reagents for non-traditional species will facilitate measurement of immune responses in wildlife (Flies et al., 2020a). When vaccines for novel pathogens affecting wild species first require development however, the challenges are more complex. Although the study described here demonstrates that DFTD vaccine research and development needs to advance, there are encouraging results from other wildlife vaccine spheres. A decade of research on koala chlamydia vaccine development has resulted in a vaccine with both protective and therapeutic effects (Waugh et al., 2020), and candidate vaccine trials for the fungal White Nose Syndrome in bats have shown promising results (Rocke et al., 2019). These examples are impressive when considering the length of time taken to develop a chlamydia vaccine for humans (Abraham et al., 2019, Lizárraga et al., 2019); and that there are no licensed fungal vaccines for human use.

In summary, the experimental DFT1 vaccination protocol used in the Stony Head trial was not protective. However, humoral and/ or cell mediated immune responses were demonstrated in the vaccinated devils that were re-trapped and this contrasts with the typical lack of immune response against DFT1. Conditions do occur under which effective anti-DFT1 immune responses are triggered, and some of these responses have resulted in immune mediated tumour regression, both natural and experimental. This information, and the results presented here where vaccine-induced immune responses lasted up to two years, suggest that DFT1 vaccine development could advance to the stage of providing protective immunity against DFT1, and that protection could have a long duration relative to the lifespan of a wild devil.

## Acknowledgments

Research support was provided by the Australian Research Council (LP130100218; LP140100508; LP180100244), and the University of Tasmania Foundation through funds raised by the Save the Tasmanian Devil Appeal and Wildcare Tasmania. We thank Narelle Phillips for assistance with the immunohistochemistry studies, and veterinary pathologists Graeme Knowles, Andrew Thompson, Andrew Davis and Russell Graydon for the tumour biopsy histopathology reports. We are grateful to the Department of Defence for access to the Stony Head military site, and to Ben and Barbara McBride for site access and field assistance.

