## Supplemental Tables S1 to S6 for "Post release immune responses of Tasmanian devils vaccinated with an experimental devil facial tumour disease vaccine"

**Table S1.** Summary of the Animal Health Laboratory* histopathology reports for the DFT1 biopsies collected from the individual devils referred to in the manuscript. Vaccinated devils, Va1 to Va8; and incumbent devils, In1 to In8.

| Devil ID | Date of tumour biopsy | Number of mitotic figures per high power field | Comments |
|---|---|---|---|
| **Va4** | Aug 2018 | Rare | Moderate anisokaryosis & karyomegaly |
| **Va5** | Sep 2017 | 2 | Minimal pleomorphism |
| **Va6** | June 2017 | Rare | Mild anisocytosis and anisokaryosis |
| | Sep 2017 | 1-3 | Mild to moderate anisokaryosis and karyomegaly, moderate pleomorphism |
| **Va7** | June 2017 | 1-2 | Mild anisocytosis and anisokaryosis |
| | Sep 2017 | 2 | Pleomorphism increased compared to June 2017 |
| | Feb 2018 | 1-2 | Mild to moderate anisokaryosis and karyomegaly |
| **In1** | Feb 2018 | 1-2 | Mild to moderate anisokaryosis and karyomegaly |
| **In2** | May 2018 | 1-2 | Mild to moderate anisokaryosis and karyomegaly |
| **In3** | May 2018 | 1-2 | Mild to moderate anisokaryosis and karyomegaly |
| **In4** | May 2018 | 1-2 | Mild to moderate anisokaryosis and karyomegaly |
| **In5** | May 2018 | 1-2 | Mild to moderate anisokaryosis and karyomegaly |
| **In6** | Feb 2018 | 1-2 | Mild to moderate anisokaryosis and karyomegaly |
| **In7** | Feb 2018 | 1-2 | Mild to moderate anisokaryosis and karyomegaly |
| | May 2018 | 1-2 | Mild to moderate anisokaryosis and karyomegaly Extensive inflammation expanding the connective tissue between nodules |
| **In8** | Feb 2018 | 1-2 | Mild to moderate anisokaryosis and karyomegaly Apoptotic tumour cells in centre of multiple nodules |

* Animal Health Laboratory, Department of Primary Industries, Parks, Water and the Environment, Prospect, Tasmania, 7250

**Table S2.** Sex, year of birth, house names and microchip numbers of the individual Tasmanian devils referred to in the manuscript**.**

| Devil identification in manuscript | Sex | Year of birth | House name | Microchip number |
|---|---|---|---|---|
| **Va1** | M | 2013 | Akaroa | 982000153632053 |
| **Va2** | M | 2014 | Askja | 982000191016491 |
| **Va3** | M | 2014 | Geysir | 982000123208479 |
| **Va4** | F | 2013 | Guernsey | 982000191016028 |
| **Va5** | M | 2013 | Macca | 982000191010136 |
| **Va6** | M | 2014 | Moffett | 982000123209291 |
| **Va7** | F | 2012 | Nutella | 982009106163346 |
| **Va8** | F | 2014 | Ventoux | 982000363283037 |
| **In1** | M | 2015 | Brimstone | 982000405800521 |
| **In2** | M | 2016 | Clay | 982000405978328 |
| **In3** | F | 2016 | Lapis | 982000363454835 |
| **In4** | M | 2016 | Mercury | 982000405977541 |
| **In5** | M | 2016 | Obsidian | 982000405977892 |
| **In6** | M | 2015 | Pumpkin | 982000405978897 |
| **In7** | F | 2016 | Ruby | 982000405800801 |
| **In8** | F | 2015 | Zuzanna | 982000363410824 |
| **In9** | F | 2013 | Tilly | 982000405949360 |

**Table S3:** Summary of immunohistochemistry analysis of DFT1 biopsies for devils trapped once, showing the average number of tumour infiltrating cells with standard deviations. The number in brackets is the number of high powered fields counted for each tumour. Note not all biopsies had enough tumour tissue to count 10 fields*. Vaccinated devils, Va4 and Va5; and incumbent devils, In1-In6, In8.

| **Devil ID** | **Tumour** | **IHC stain** | | |
|---|---|---|---|---|
| | | **CD3** | **MHC-II** | **PD-1** |
| **Va4** | **T1** | 11 ± 5 (3) | 17 ± 9 (2) | 9 (1) |
| **Va5** | **T1** | 15 ± 6 (9) | 20 ± 8 (8) | 2 ± 2 (8) |
| | **T2** | 14 ± 11 (6) | 16 ± 6 (4) | 4 ± 4 (4) |
| **In1** | **T2** | 2 ± 1 (10) | 5 ± 3 (8) | 0 (5) |
| **In2** | **T1** | 3 ± 1 (8) | 4 ± 2 (7) | 0 (5) |
| | **T4** | 1 ± 1 (6) | 7 ± 2 (6) | NA |
| **In3** | **T1** | 0 (8) | 2 ± 2 (7) | 0 (5) |
| **In4** | **T1** | 1 ± 1 (8) | 1 ± 1 (7) | 0 (5) |
| **In5** | **T1** | 0 (8) | 3 ± 3 (8) | 0 (5) |
| **In6** | **T2** | 2 ± 2 (7) | 6 ± 2 (6) | 3 ± 2 (6) |
| **In8** | **T1** | 2 ± 1 (8) | 10 ± 3 (7) | 0 (5) |

NA = not available

*Method according to (Zhang et al., 2003)

**Table S4:** Summary of the immunohistochemistry analysis of tumours biopsied from vaccinated devils, Va6 and Va7, and incumbent devil, In7, on sequential monitoring trips, showing the degree of immune cell infiltration in the tumour biopsies.

| **Biopsy date** | | **June 2017** | | | **Sep 2017** | | | **Feb 2018** | | | **May 2018** | | |
|---|---|---|---|---|---|---|---|---|---|---|---|---|---|
| **Immunohistochemistry stain** | | **CD3** | **MHCII** | **PD1** | **CD3** | **MHCII** | **PD1** | **CD3** | **MHCII** | **PD1** | **CD3** | **MHCII** | **PD1** |
| **Va6** | T1 | 0 | + | 0 | + | + | 0 | Not trapped | | | Not trapped | | |
| **Va7** | T1 | + | + | 0 | ++ | ++ | 0 | +++ | +++ | ++ | Not trapped | | |
| | T3 | NA | NA | NA | NA | NA | NA | ++ | ++ | + | | | |
| **In7** | T1 | Not trapped | | | Not trapped | | | + | + | 0 | ++ | ++ | ++ |

T1: tumour 1; T3: tumour 3

NA: not applicable

Intratumoural immune cells graded as 0, +, ++, or +++ (0, 1-5, 6 to 19, or ≥20 cells per high-power field, respectively) (Zhang et al., 2003)

**Table S5**. Summary of immunohistochemistry analysis of tumours biopsied from vaccinated devils, Va6 and Va7, and incumbent devil, In7, on sequential monitoring trips showing the average number of infiltrating tumour cells with standard deviations. The number in brackets is the number of high powered fields counted for each tumour*. Note not all biopsies had enough tumour tissue to count 10 fields.

| **Biopsy date** | | **06/17** | | | **09/17** | | | **02/18** | | | **05/18** | | |
|---|---|---|---|---|---|---|---|---|---|---|---|---|---|
| **IHC stain** | | **CD3** | **MHCII** | **PD1** | **CD3** | **MHCII** | **PD1** | **CD3** | **MHCII** | **PD1** | **CD3** | **MHCII** | **PD1** |
| **Va6** | **T1** | 0 (6) | 3 ± 2 (5) | NA | 1 ± 1 (8) | 5 ± 5 (6) | 0 (5) | Not trapped again | | | | | |
| **Va7** | **T1** | 2 ± 2 (10) | 5 ± 2 (10) | NA | 10 ± 6 (10) | 13 ± 6 (8) | NA | 31 ± 4 (5) | 33 ± 9 (6) | 15± 1 (3) | Not trapped again | | |
| | **T3** | NA | NA | NA | NA | NA | NA | 8 ± 3 (6) | 13 ± 3 (5) | 4 ± 2 (3) | | | |
| **In7** | **T1** | Not trapped until 02/18 | | | | | | 1 ± 1 (8) | 5 ± 4 (7) | NA | 12 ± 10 (8) | 19 ± 5 (9) | 8 ± 2 (5) |

NA = not available

*Method according to (Zhang et al., 2003)

**Table S6.** Summary of the serum antibody responses against MHC-I$^{-ve}$ and MHC-I$^{+ve}$ DFT1 cells of all vaccinated devils (Va1 to Va8), and of the incumbent devils that had a positive result (In1, In8, In9). Results are shown for the serum samples collected on the last date the devils were trapped post-release. Antibody responses were classified as negative "-" (MFIR < 1.5), or positive "+" (MFIR ≥ 1.5) (Pye et al., 2018).

| Devil ID | Sex; age (in years) at time of sample collection | DFT1 status, date of DFT1 diagnosis (or, if healthy, date last trapped) | Serum antibodies against MHC-I $^{-ve}$ DFT1 cells | Serum antibodies against MHC-I $^{+ve}$ DFT1 cells |
|---|---|---|---|---|
| **Va1** | M 4y | Healthy 09/17 | + | + |
| **Va2** | M 3y | DFT1 07/17 | + | + |
| **Va3** | M 3 y | Healthy 09/17 | + | + |
| **Va4** | F 5 y | DFT1 08/18 | - | + |
| **Va5** | M 4y | DFT1 09/17 | + | + |
| **Va6** | M 3y | DFT1 07/17 | + | + |
| **Va7** | F 5y | DFT1 07/17 | + | + |
| **Va8** | F 3y | DFT1 Trapped healthy 07/17. Found dead with DFT1 08/18 | + | + |
| **In1** | M 3y | DFT1 02/18 | + | + |
| **In8** | F 3y | DFT1 02/18 | - | + |
| **In9** | F 5y | Healthy 03/19 | - | + |

**Table S7** Summary of the age, sex and DFT1 status of the 76 incumbent devils that were tested for serum antibody against DFT1 cells. Three devils (In1, In8, In9) out of the 76 devils were positive for serum antibody and these are referred to in Tables S2 and S8

| **Age** | **Healthy** | | **DFT1** | |
|---|---|---|---|---|
| | **Male** | **Female** | **Male** | **Female** |
| **Subadult (<2yo)** | 26 | 26 | 2 | 2 |
| **Adult (≥2yo)** | 4 | 6 | 4 | 6 |
